# *Teladorsagia circumcincta* 1,6 bisphosphate aldolase: molecular and biochemical characterisation, structure analysis and recognition by immune hosts

**DOI:** 10.1101/2020.07.13.201335

**Authors:** S. Umair, C.L.G. Bouchet, N. Palevich, J.S. Knight, H.V. Simpson

## Abstract

A 1095 bp full length cDNA encoding *Teladorsagia circumcincta* aldolase (*Tci*ALDO) was cloned, expressed in *Escherichia coli,* the recombinant protein purified and its kinetic properties determined. A phylogenetic tree was constructed using helminth aldolase sequences. The predicted protein consisted of 365 amino acids and was present as a single band of about 44 kDa on SDS-PAGE. Multiple alignments of the protein sequence of *Tci*ALDO with homologues from other helminths showed that the greatest similarity (93%) to the aldolases of *Haemonchus contortus* and *Dictyocaulus viviparus*, 82-86% similarity to the other nematode sequences and 68-71% similarity to cestode and trematode enzymes. Substrate binding sites and conserved regions were identified and were completely conserved in other homologues. At 25 °C, the optimum pH for *Tci*ALDO activity was pH 7.5, the V_max_ was 432 ± 23 nmoles.min^−1^.mg^−1^ protein and the apparent K_m_ for the substrate fructose 1,6-bisphosphate was 0.24 ± 0.01 μM (mean ± SEM, n = 3). Antibodies in both serum and saliva from field-immune, but not nematode-naïve, sheep recognised recombinant *Tci*ALDO in enzyme-linked immunosorbent assays. The recognition of the recombinant protein by antibodies generated by exposure of sheep to native aldolase indicates similar antigenicity of the two proteins.

## 1. Introduction

Fructose 1,6-bisphosphate aldolase (FBA) (EC 4.1.2.13) catalyses the reversible reaction that splits fructose 1,6-bisphosphate into the triose phosphates dihydroxyacetone phosphate (DHAP) and glyceraldehyde 3-phosphate (G3P). The forward reaction occurs during glycolysis and the reverse reaction forms fructose 1,6-bisphosphate during gluconeogenesis. FBA enzymes belong to two classes depending on the mechanism of the reaction: class I, which form covalent Schiff-base conjugates with a conserved lysine, are present mainly in higher eukaryotes and a few bacteria, whereas class II require a divalent metal ion as cofactor for enzymatic activity and are found principally in bacteria, algae and fungi (Grazi et al., 1962; Marsh and Lebherz, 1992; Gefflaut et al., 1996; Sánchez et al., 2002; Tittmann). Thus, class I and II enzymes can be distinguished by inhibition of the latter by ethylene diamine tetraacetic acid (EDTA). There are three isoforms of vertebrate FBA: aldolase A which is principally expressed in muscle, aldolase B in liver and aldolase C in brain (Lebherz and Rutter, 1969).

The genes encoding FBAs have been sequenced from the free-living nematode *Caenorhabditis elegans* (Inoue et al., 1997), the animal-parasitic *Haemonchus contortus* (Yan et al., 2013), the plant-parasitic *Heterodera glycines* and *Globodera rostochiensis* (Kovaleva et al., 2005), as well as other helminths, including *Schistosoma mansoni* (El-Dabaa et al., 1998) *Echinococcus granolosus* (Lorenzatto et al., 2012), *Clonorchis sinensis* (Li et al., 2014), *Schistosoma japonicum* (Hu et al., 2015) and *Opisthorchis viverrini* (Prompipak et al., 2017). As nematode FBAs were shown to have some structural properties similar to vertebrate FBA A, but catalytic properties more like those of FBA C, aldolases were suggested to be the products of primordial genes from which vertebrate FBA genes have evolved (Reznick and Gershon, 1977a). Subsequent genetic studies have shown that in *C. elegans* there are two isozymes encoded by different genes, one of which has similar kinetic properties to vertebrate aldolase C and the other broader substrate specificity in addition to fructose 1,6-bisphosphate (Inoue et al., 1997), which could explain the earlier conclusions about nematode aldolases being evolved by duplication and diversification.

The kinetic properties of FBA enzymes are generally similar, such as the typical temperature and pH optima of 40°C and pH 7.5 respectively of the *H. contortus* enzyme (Yan et al., 2013). The reported K_m_ of purified nematode aldolases varied between species and even within studies, e.g. the aldolase in homogenates of *H. glycines* had a lower activity than either the *C. elegans* or *Panagrellus redivivus* enzymes (Reznick and Gershon, 1977b) because of there might be a mixture of different isozymes. Enzyme activity declined with age in the free-living *Turbatrix aceti* (Zeelon et al., 1973). Parasitic helminths may have more active enzymes than their hosts, as seen for *S. japonicum* FBA, which had a lower apparent K_m_ of 0.06 μM and higher activity than that of human FBA A (Hu et al., 2015).

Aldolase, like many other glycolytic enzymes, has both intra- and extra-cellular further activities in parasites in addition to its enzymatic function (Gόmez-Arreaza et al., 2014); these include plasminogen binding (Ramajo-Hernández et al., 2007) and immunomodulation (Marques et al., 2008). It is released into the extracellular environment and can be detected in excretory/secretory (ES) products (Ramajo-Hernández et al., 2007; Guillou et al., 2007; Morassutti et al., 2012) and has been also located in the tegument of adult *Schistosoma bovis* (Ramajo-Hernández et al., 2007) and *S. mansoni* (El-Dabaa et al., 1998). FBA is antigenic: the protein from *Dirofilaria immitis* is recognised by the sera of infected dogs (Sassi et al., 2014); the serum of *Onchocerca volvulus* infected and putatively immune humans recognised aldolase (McCarthy et al., 2002), *Angiostrongylus cantonensis* (Morassutti et al., 2012) or *S. japonicum* (Peng et al., 2009) and the *Teladorsagia circumcincta* L3 protein reacted with ovine IgA in the gastric lymph of previously-infected sheep (Ellis et al., 2014).

In the present study, the cDNA encoding *T. circumcincta* aldolase (*Tci*ALDO) was cloned, expressed in *Escherichia coli*, the recombinant protein was purified and some kinetic properties determined. Enzyme-linked immunosorbent assays (ELISAs) were performed to determine if the enzyme was recognised by saliva and serum from sheep previously exposed to nematode parasites in the field. To gain insight into the substrate specificity of *Tci*ALDO, we modelled the crystal and tertiary structures using the full-length amino acid sequence of *Tci*ALDO.

## 2. Material and methods

All chemicals were purchased from the Sigma Chemical Co. (Mo, USA) unless stated otherwise. Use of experimental animals for culturing and harvesting adult worms for RNA extraction has been approved by the AgResearch Grasslands Animal Ethics Committee (protocol #13052).

### 2.1. Parasites

Pure cultures of *T. circumcincta* were maintained in the laboratory by regular passage through sheep. Adult worms were recovered from the abomasa of infected sheep as described previously (Umair et al., 2013). Briefly, abomasal contents were mixed 2:1 with 3% agar and the solidified agar blocks incubated at 37 °C in a saline bath. Clumps of parasites were collected from the saline soon after emergence and frozen in Eppendorff tubes at −80 °C for molecular biology procedures.

### 2.2. RNA isolation and cDNA synthesis

Adult *T. circumcincta* (50-100 μl packed volume) in 1 ml Trizol (Life Technologies) were ground to a fine powder in a mortar under liquid N_2_ and RNA extracted according to the manufacturer’s instructions. The quality and concentration of the RNA was assessed and first strand cDNA synthesised from 1 ug using the iScript Select cDNA Synthesis Kit (Bio-Rad) and a 1:1 mixture of Oligo (dT) _20_ and hexamer primers.

### 2.3. *Cloning and expression of* T. circumcincta *recombinant* Tci*ALDO in* E. coli

A partial *T. circumcincta* ALDO sequence TDC00486 (NEMBASE) containing the 5’ end was used and the 3’ of *Tci*ALDO cDNA was obtained by 3’ RACE (SMARTer RACE cDNA Amplification Kit, Clontech) using *T. circumcincta* adult RNA, as outlined by the manufacturer. The full length *Tci*ALDO cDNA was amplified from this cDNA in a PCR containing the oligonucleotide primers Tci aldo_FL-F1 (5’-CACCATGGCTTCCTACTCGCAGTA-3’) and Tci aldo_FL-R1 (5’-TCAATAGGCATGATTAGCCAC-3’). The full-length gene was then transformed into TOP10 cells and subsequently cloned into the expression vector Champion pET100 Directional TOPO (ThermoFisher Scientific) and transformed into *E. coli* One shot BL21 (DE3) according to the manufacturer’s instruction. The construction was sequenced to check its integrity.

*E. coli* strain BL21 (DE3) transformed with pET 100 using a NH_2_ tag *Tci*ALDO was grown in 10 ml Luria Broth (LB) supplemented with 100 μg/ml ampicillin for 16 h at 30 °C and 250 rpm. The culture was diluted 20-fold in LB with 100 μg/ml ampicillin and 1% glucose and grown to OD_600_ of 0.6-0.8 at 30 °C and 250 rpm. Isopropyl-1-thio-β-D-galactopyranoside (IPTG) was added to a final concentration of 1 mM and the culture grown at 30°C and 250 rpm for an additional 16. Bacteria were harvested by centrifugation and the soluble extract was obtained by incubating harvested cells in CelLyticB (5ml/g wet cell paste), 0.2 mg/ml Lysozyme, 50U/ml Benzonase and protease inhibitors at room temperature for 10 min. The crude lysate was centrifuged at 15,000x*g* at 4 °C for 20 min to remove all the cell debris and the supernatant collected and filtered through a 0.45 μm filter.

### 2.4. Purification

Purified recombinant poly-histidine protein was obtained by FPLC under native conditions using a Ni-NTA column (Qiagen), and a Biologic DUO-FLOW BIO-RAD chromatography system (Bio-Rad, USA). Sodium bi-phosphate buffer was used as an equilibration buffer, sodium bi-phosphate containing 20 mM imidazole as the wash buffer and sodium bi-phosphate containing 500 mM imidazole as elution buffer. The protein was dialysed overnight following the elution and the concentration was determined by the Nanodrop (Thermofisher Scientific) using the A280 nm assay with extinction coefficient (34755 M^−1^cm^−1^) and molecular weight (43.8 KDa).

### 2.5. Bioinformatics

Alignment of protein sequences was performed using the Muscle alignment option in Geneious Prime (Biomatters Ltd) with the Blosum 62 similarity matrix used to determine similarity to *H. contortus* and other helminth aldolases. The predicted tetramer structure of *Tci*ALDO was constructed using SWISS-MODEL, a fully automated protein structure homology-modelling server with default parameters.

### 2.6. Gel electrophoresis

SDS-PAGE was performed using NuPAGE Novex 4-12% Bis-Tris gels according to the instructions of the manufacturer (Invitrogen). Gels were stained with SimplyBlue safe stain (Life technologies). A western blot was also performed using a monoclonal anti-poly histidine-peroxidase antibody. Blots were blocked, incubated overnight in 1:2000 antibody in buffer (4% skim milk powder in Tris-buffered saline and 0.1% Tween-20) at room temperature and developed to detect His-tagged recombinant protein.

### 2.7. Tci*ALDO activity (E.C. 4.2.1.11)*

The enzyme activity of *Tci*ALDO was measured at 30 °C in a coupled assay with reversible conversion of fructose 1,6-bisphosphate to glyceraldehyde 3-phosphate and dihydroxyacetone phosphate using Sigma aldolase kit (Catalogue # MAK223). The NADH production was measured colorimetrically at 450 nm. The final reaction mixture (100 ul) contained assay buffer, enzyme mix, enzyme developer, recombinant protein (50 μg) and the substrate. NADH standards and the blank were set up as described by the manufacturer.

1. The optimum pH was determined with a substrate concentration of 0.5 mM fructose 1,6-bisphosphate with pH range 6 to 9. Subsequent assays were carried out at pH 7.5.
2. The apparent K_m_ for fructose 1,6-bisphosphate was determined in reaction mixtures containing 0-5 mM fructose 1,6-bisphosphate.
3. The effects of EDTA as potential activators/inhibitors on recombinant *TciALDO* with substrate concentrations of 0.5 mM fructose 1,6-bisphosphate and 10 mM EDTA.

### 2.8. ELISA

Serum and saliva samples taken from parasite-naive and parasite-exposed sheep were tested for the presence of antibodies that react with the recombinant *Tci*ALDO by ELISA. Serum and saliva samples were collected from 18 male Romney lambs 6-7 months old and previously exposed to multiple species of parasites including *H. contortus* and *T. circumcincta*. Lambs had developed immunity against *T. circumcincta* infection and pooled serum and saliva samples were used for ELISA. Plates were coated with 5 μg/ml *Tci*ALDO onto ELISA plates (Maxisorp, Thermofisher Scientific) overnight at 4 °C. Plates were washed and free binding sites were then blocked with Superblock (Thermofisher Scientific) at room temperature for 30 min and then incubated with serial dilutions (200- to 12800-fold for serum) or (20- to 320-fold for saliva) in ELISA buffer for 2h at room temperature. Bound serum immunoglobulins were then detected with rabbit anti-sheep IgG-HRP diluted 1:4000 by incubation at 37°C for 2h and colour development with 3,3’,5,5’ tetramethylbenzidine (TMB). Saliva IgA was detected with rabbit anti-sheep IgA-HRP diluted, incubated and colour was developed as described for serum IgG.

### 2.9. Protein modelling and structural analysis of *Tci*ALDO

The structural model of *Tci*ALDO was constructed by submitting the amino acid sequence obtained as described above, to the I-TASSER server (Yang et al., 2015). From the best five models obtained, a model was selected with a C-score of −0.11, a TM value of 0.70 ± 0.12. TM-score is a metric for measuring the similarity of two protein structures, or a global fold similarity between the generated model and the structure it was based on. Scores higher than 0.5 assumes the parent structure and modelled protein share the same fold while below 0.17 suggests a random nature to the produced model (Zhang and Skolnick, 2004). C-score is a confidence score for estimating the quality of predicted models by I-TASSER. It is calculated based on the significance of threading template alignments and the convergence parameters of the structure assembly simulations. C-score is typically in the range of −5 to 2, where a C-score of higher value signifies a model with a high confidence and vice-versa. The structural model with highest C-score was further validated using Procheck (Laskowski et al., 1996) and ProSA-web (Wiederstein and Sippl, 2007). The substrate Binding domain was identified and active site residues were deduced and pictured using PyMol (Schrodinger, 2010).

### 2.10. Data analysis

Replicate data are presented as mean ± SEM. Graph Prism v5 was used to plot kinetic data and estimate K_m_ and V_max_. The kinetic data were analysed using the non-linear fit function of Graph Prism and the best fit shown to be a one-site binding hyperbola.

## 3 Results

### 3.1. *Tci*ALDO gene sequence

The 1,095 bp full length *T. circumcincta* cDNA sequence, amplified from adult *T. circumcincta* cDNA, has been deposited in Genbank as Accession No **KX452943**. The predicted protein consisted of 365 amino acids (Fig. 1). A multiple alignment, using Alignment Geneious Prime, of the protein sequences of *Tci*ALDO with homologues from *H. contortus, C. elegans, Caenorhabditis briggsae, Ancylostoma ceylanicum, C. sinensis, E. granulosus, Necator americanus*, *S. japonicum, O. viverrini* and *H. glycines* is shown in Fig. 1. Substrate binding sites and conserved regions in other homologues were identified and shown to be completely conserved in *Tci*ALDO.

**Fig. 1.**
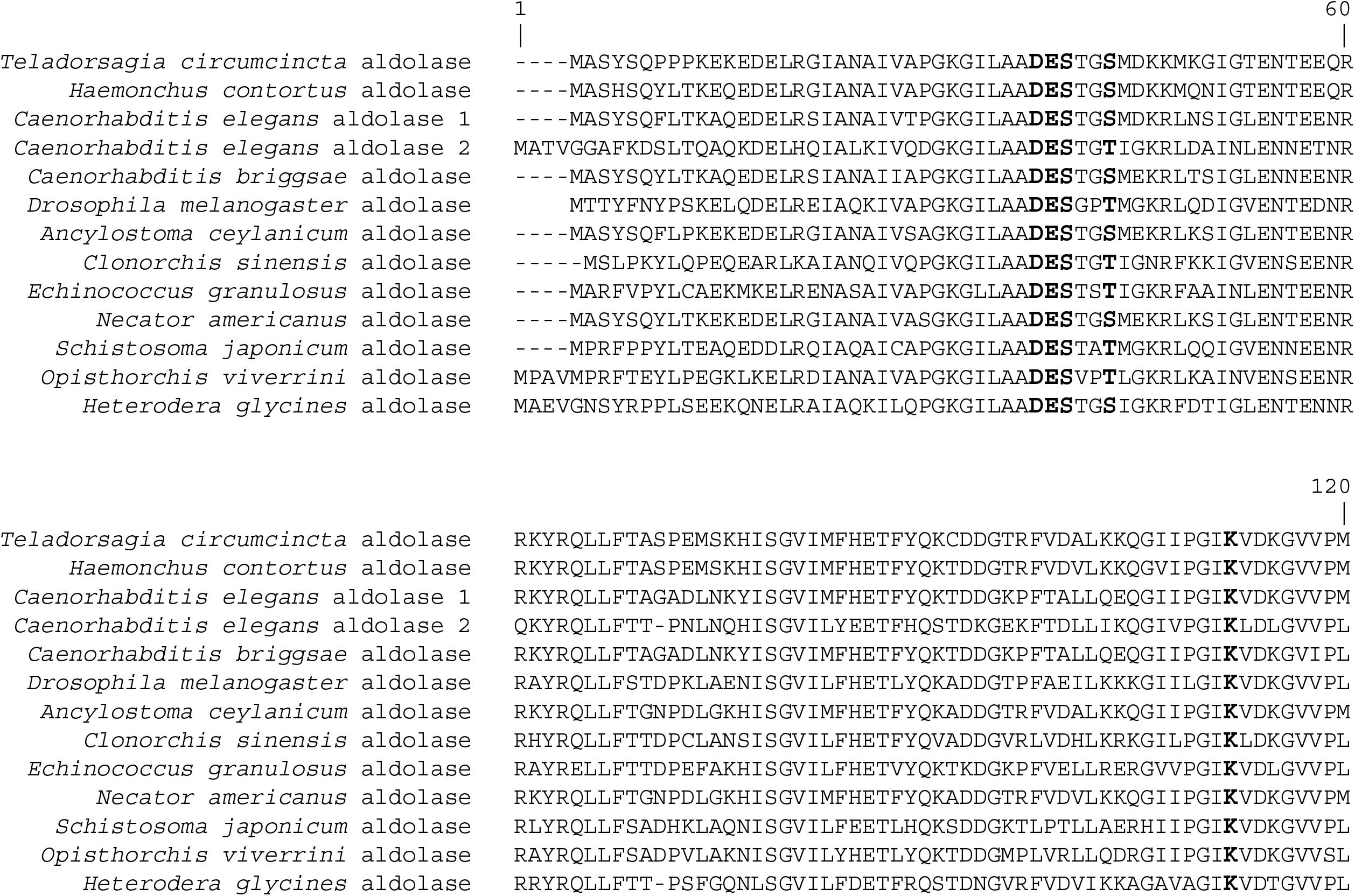
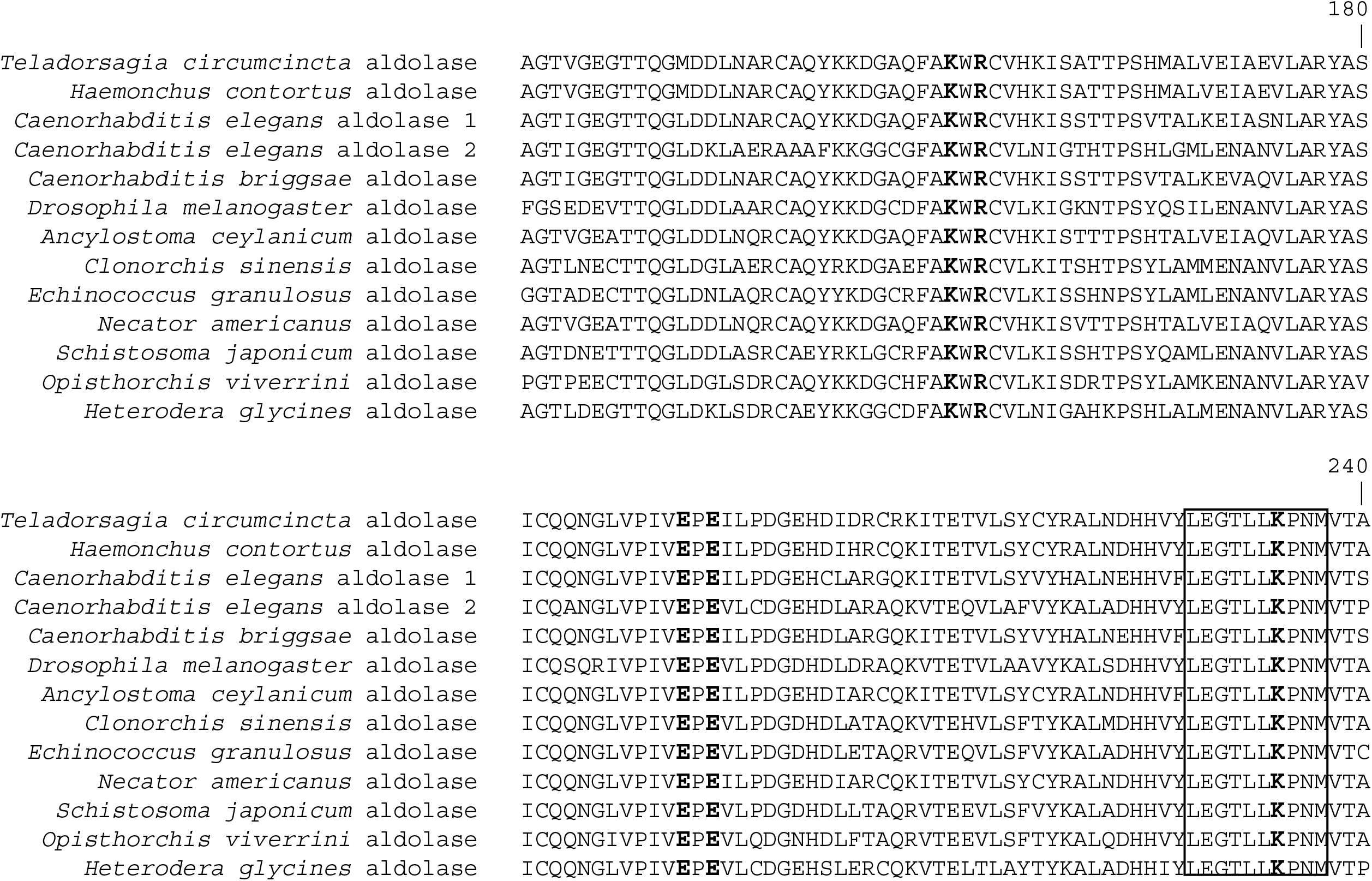
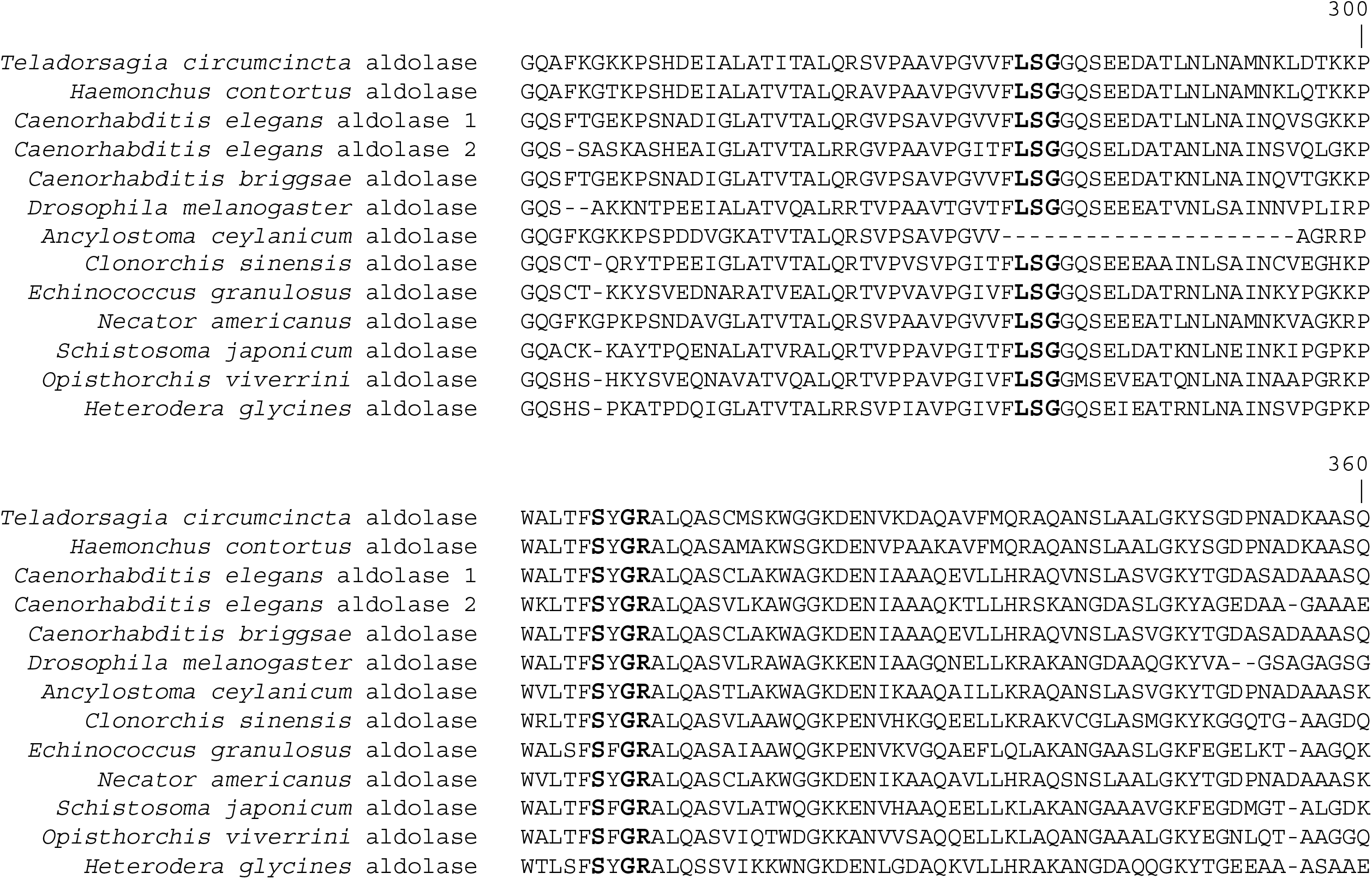
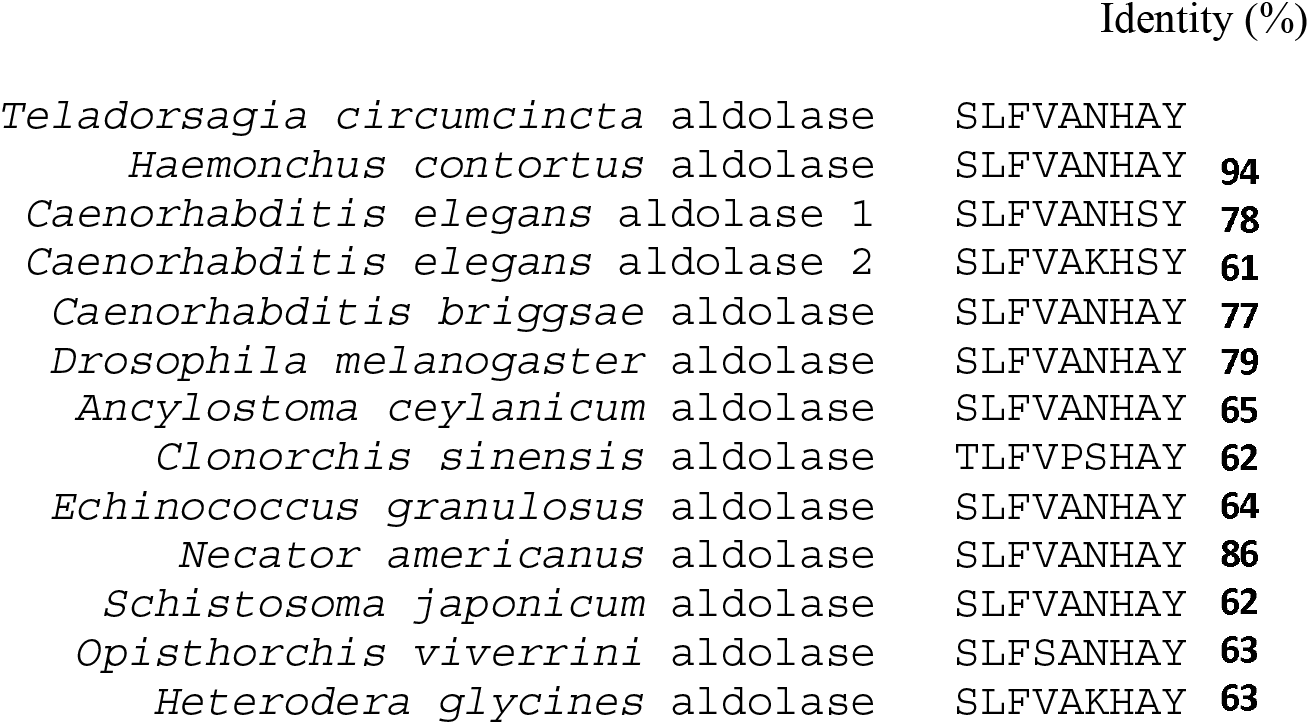
Multiple sequence alignment of aldolases from *T. circumcincta* (GI: **KX452943),***H. contortus* (GI: **ADT61995**), *Caenorhabditis elegans* (GI: **CAB03291**), *Caenorhabditis elegans* aldolase 2 (GI: **CCD65997**), *Caenorhabditis briggsae* (GI: **XP002643138**), *Ancylostoma ceylanicum* (GI: **EPB73313**), *Clonorchis sinensis* (GI: **GAA50927**), *Echinococcus granulosus* (GI: **EUB64508**), *Necator americanus* (GI: **XP013291330**), *Schistosoma japonicum* (GI: **CAX78614**), *Opisthorchis viverrini* (GI: **OON18662**) and *Heterodera glycines* (GI: **AAG47838**), homologues. Amino acid residues indicated in the marked box are essential to the aldolase activity.

To identify the active site as well as infer both functional and structural characteristics, we modelled the 3D structure of *Tci*ALDO using the I-TASSER server (Fig. 2). The C-score of the best five models were less than −2.9, expected TM Score was < 0.7 and normalized z-scores were less than 7.93. the ITASSER modelled protein produced was similar to the parent molecule of with a C-score of −0.11 and a TM value of 0.70 all within acceptable ranges.

**Fig. 2.**
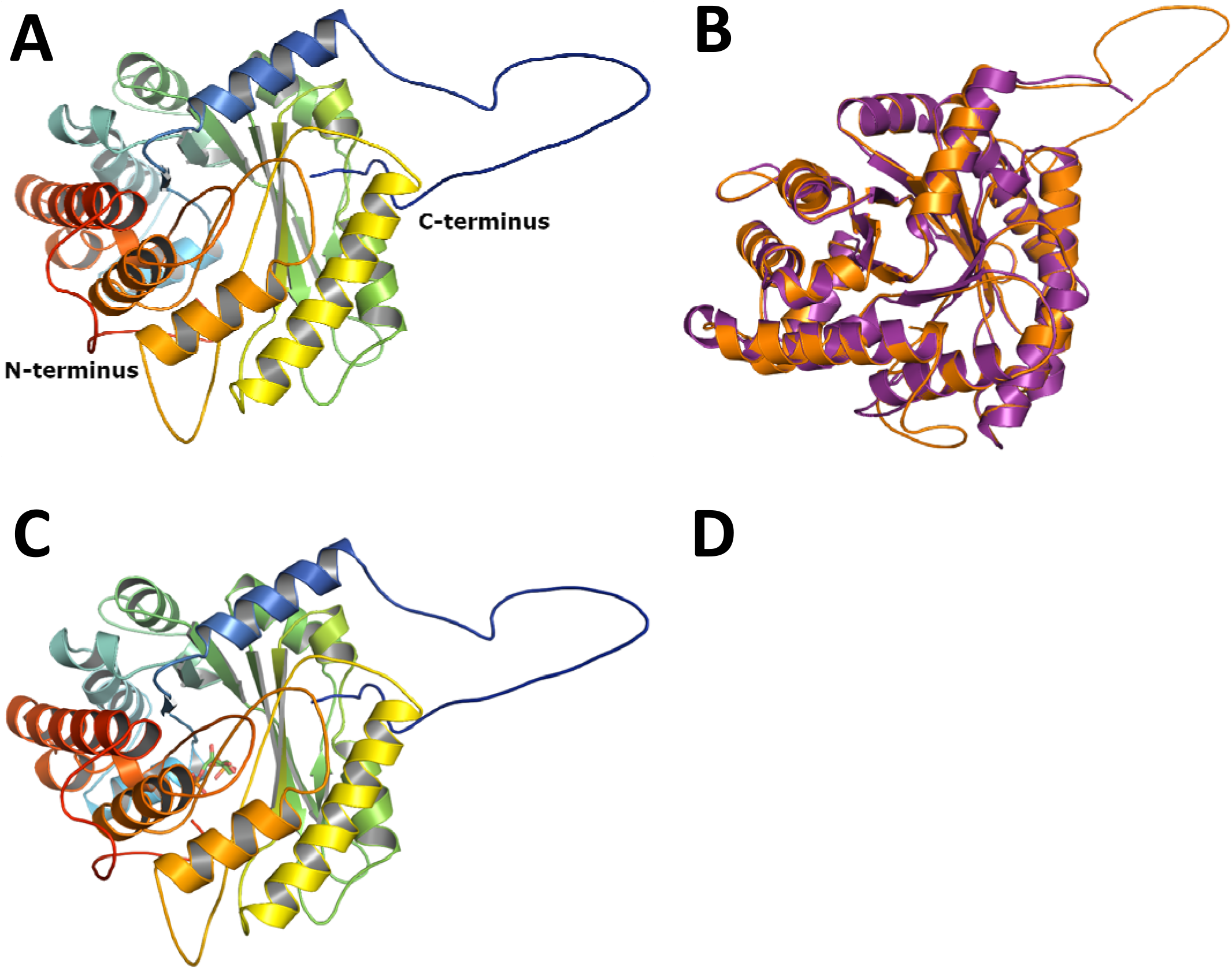
(A) The predicted tertiary structure of *Tci*ALDO monomer (B) Superposition of the predicted tertiary structure of *Tci*ALDO from *T. circumcincta* (orange) and 3TU9 (purple). 3TU9 is the repository code for the coordinates of rabbit muscle aldolase stored at the protein data bank (https://www.rcsb.org/structure/3TU9), and was the template for our model (C) Location of the active site within *Tci*ALDO. (D) The active site of *Tci*ALDO (in green) within 4Å of the superimposed 2FP (in blue). 2FP is abbreviation for “1,6-fructose diphosphate” is also stored at the protein data bank (https://www.rcsb.org/ligand/2FP), and was the ligand modelled for our structure.

Shown in Fig. 2 is the protein structure for *Tci*ALDO, superimposed best structural model corresponding to the monomer of 3TU9 (Mabiala-Bassiloua et al., 2011), as well as the 1,6-fructose diphosphate ligand (2FP) binding site and catalytic and active site residues that fall within 4 Å of the substrate (Ala-68, Ser-75, Ser-72, Glu-71, Asp-70, Lys-144, Lys-183, Arg-185, Glu-224, Lys-266, Leu-308, Gly-310, Ser-309, Tyr-339, Arg-341 and Gly-340). The lysine at position 230 is the residue where Schiff base intermediates are formed.

### 3.2. Recombinant protein expression

A number of varying conditions were used in the trial expression and based on which maximal production of functional recombinant ALDO was obtained in the *E. coli* strain BL21 (DE3) when expression was induced with 1 mM IPTG at 30 °C for 16. The purified N-terminal His recombinant *Tci*ALDO protein appeared as a single band of about 44 kDa on SDS-PAGE (Fig. 3). The presence of a His-Tagged recombinant protein as well as a high purity of the recombinant protein were confirmed by Western blotting.

**Fig. 3.**
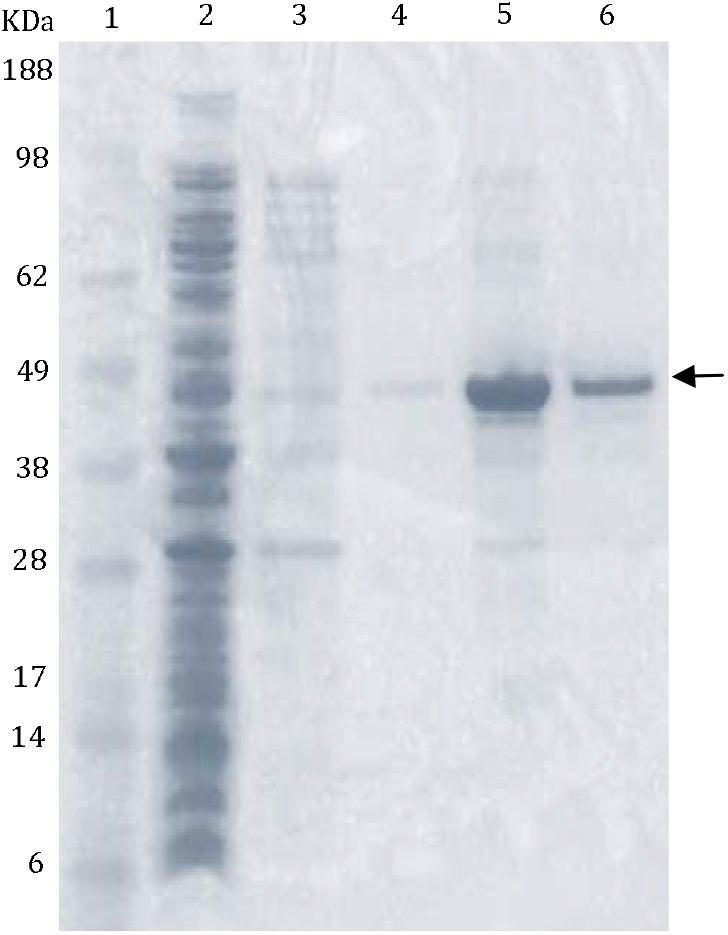
SDS-PAGE of recombinant *Tci*ALDO. Lane1: standards; lane 2: unbound; lane 3: wash 1; lane 4: wash 2; lane 5: elution 1; lane 6: elution 2; lane 7: standards.

### 3.3. Enzyme assays

The optimum pH for recombinant *Tci*ALDO activity at 25°C was 7.5 (Fig. 4). The apparent K_m_ for fructose 1,6-bisphosphate was 0.24 ± 0.01 μM and the V_max_ was 432 nmoles min^−1^ mg^−1^ protein (mean ± SEM, n=3) (Fig. 5). The Hill Coefficient was calculated to be 1.70.

**Fig. 4.**
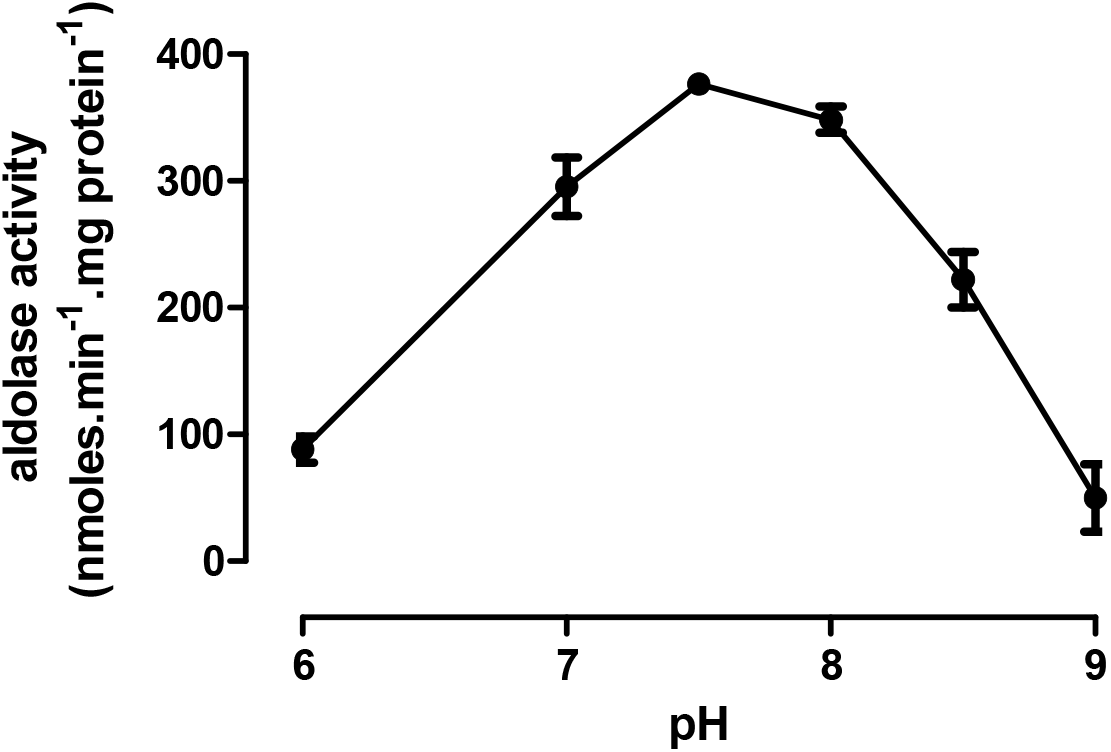
Effects of pH on the activity of recombinant *Tci*ALDO at 25 °C (mean ± SEM, *n* = 3). Enzyme activity was estimated from the rate of NADH production, which was measured colorimetrically at 450 nm.

**Fig. 5.**
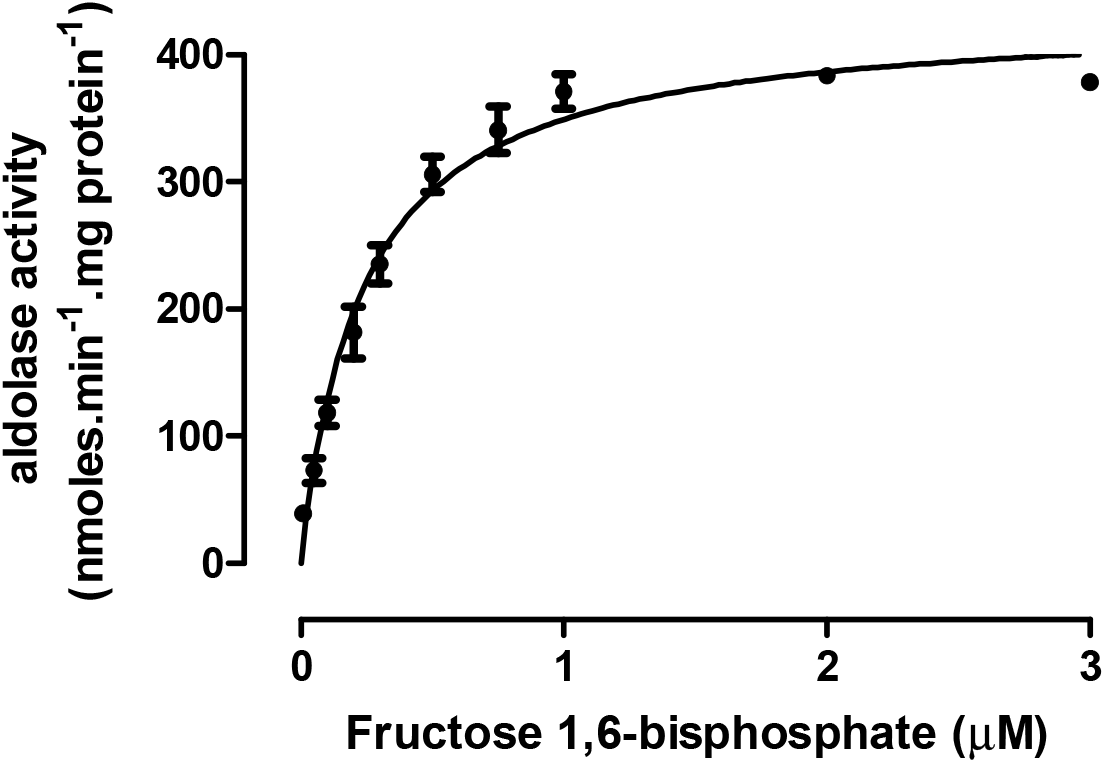
Effects of varying substrate concentration on the activity of recombinant *Tci*ALDO at 25 °C (mean ± SEM, *n* = 3). Enzyme activity was estimated from the rate of NADH production, which was measured colorimetrically at 450 nm.

### 3.4. Host recognition

Recombinant *Tci*ALDO was recognised in an ELISA by antibodies in both serum and saliva which had been collected from sheep exposed to nematodes in the field (Fig. 6). There was no antibody detection when serum or saliva samples from parasite-naïve animals was used.

**Fig. 6.**
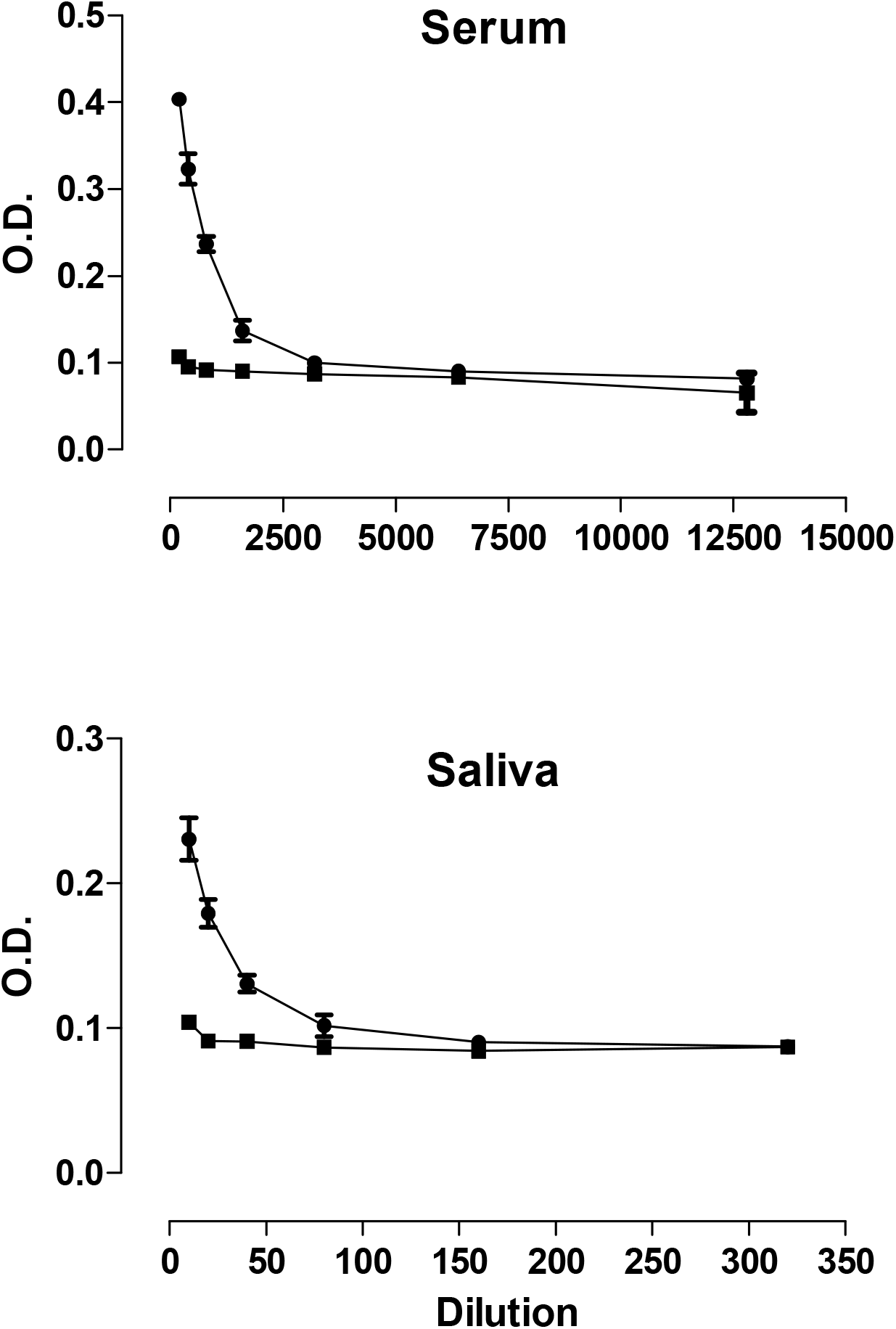
Recognition of *Tci*ALDO by serially diluted serum (IgG) (top) or saliva (IgA) (bottom) (•) from parasite exposed animals, but not by serum or saliva (■) from parasite-naïve animals.

## 4. Discussion

A 1095 bp full length cDNA sequence encoding *T. circumcincta* aldolase (*Tci*ALDO) was amplified from adult *T. circumcincta* cDNA, cloned and expressed in *E. coli*. The 365 amino acid *Tci*ALDO protein expressed in *E. coli* was typical of aldolase monomers of many species and had 77-94% similarity to the aldolase of other nematodes and 62-64 % similarity to that of cestodes and trematodes (Fig. 1).

The 3D-structure as well as binding and catalytic sites have been determined for a wide range of FBAs and are known to be highly conserved (Fig. 2) (Gefflaut et al., 1996; Dalby et al., 1999; Roslan et al., 2017). This was also true for other helminth homologues (Fig. 1), although there were minor differences in the trematode and cestode aldolase sequences, in which serine was replaced by threonine at amino acid 38.

The kinetic properties of the recombinant *Tci*ALDO were generally similar to those of enzymes of other species. The optimum pH for *Tci*ALDO activity at 25 °C was pH 7.5 (Fig. 4), similar to that for the aldolase of *H. contortus* (Yan et al., 2013). The enzyme was very active at 25 °C (V_max_ 432 ± 23 nmoles.min^−1^.mg protein^−1^), which is below its optimal temperature, assuming that it is similar to the 40-45°C of the closely related aldolases of *H. contortus* (Yan et al., 2013) and *C. sinensis* (Li et al., 2014). *Tci*ALDO activity was unchanged by the addition of 10 mM EDTA, indicating that the enzyme was class I and not class II aldolase, which is strongly inhibited by EDTA (Grazi et al., 1962; Gefflaut et al., 1996). The apparent K_m_ of *Tci*ALDO for the substrate fructose 1,6-bisphosphate was 0.2 ± 0.01 μM (Fig. 5). This is higher than the 0.06 μM reported for recombinant *S. japonicum* aldolase (Hu et al., 2015), but lower than the very variable values reported for the partially purified *Ascaris suum* (Kochman and Kwiatkowska, 1972) or *H. contortus* aldolases (Rhodes, 1972). Kinetic properties may be more accurately reflected by recombinant enzymes than purified proteins and suggest that parasitic helminths may have more active enzymes than their hosts (Hu et al., 2015).

Native *T. circumcincta* aldolase is antigenic and antibodies in both serum and saliva from field-immune, but not nematode-naïve, sheep recognised recombinant *Tci*ALDO in an ELISA (Fig. 6). Ovine IgA in the gastric lymph of previously-infected sheep reacted with aldolase in L3 *T. circumcincta* (Ellis et al., 2014). The antigenicity of aldolase has been exploited in detecting previous exposure of water buffaloes to *S. japonicum* (Peng et al., 2008), dogs to *D. immitis* (Sassi et al., 2014) or humans to *O. volvulus* (McCarthy et al., 2002), *A. cantonensis* (Morassutti et al., 2012) or *S. japonicum* (Peng et al., 2009). Aldolase, like many other glycolytic enzymes, has both intra- and extra-cellular activities in pathogens in addition to its enzymatic function (Gόmez-Arreaza et al., 2014), which have been suggested to facilitate their establishment in the host. These appear to be essential to successful establishment of many pathogens, including helminths. This is supported by the protection against infection induced by vaccination of mice with the aldolase of *S. mansoni* (Marques et al., 2008) or fish with the aldolase of several pathogenic bacteria (Sun et al., 2015).

## Acknowledgments

The authors would like to thank Drs Axel Heiser and Sandeep Gupta for critically reviewing the manuscript. The financial support of AGMARDT (Grant No. P14003) is gratefully acknowledged.

## References

Dalby, A., Dauter, Z., Littlechild, J.A., 1999. Crystal structure of human muscle aldolase complexed with fructose-1,6-bisphosphate: Mechanistic implications. Protein Science 8, 291–297.

El-Dabaa, E., Mei, H., El-Sayed, A., Karim, A.M., Eldesoky, H.M., Fahim, F.A., LoVerde, P.T., Saber, M.A., 1998. Cloning and characterization of *Schistosoma mansoni* fructose-1,6-bisphosphate aldolase isoenzyme. Journal of Parasitology 84, 954–960.

Ellis, S., Matthews, J.B., Shaw, D.J., Hamish, S.P., McWilliam, E.G., Inglis, N.F., Nisbet, A.J., 2014. Ovine IgA-reactive proteins from *Teladorsagia circumcincta* infective larvae. International Journal for Parasitology 44, 743–750.

Grazi, E., Rowley, P.T., Cheng, T., Tchola, O.L., 1962. The mechanism of action of aldolases III. Schiff base formation with lysine. Biochemistry and Biophysics Research Communication 9, 48–53.

Gefflaut, T., Blonski, C., Perie, J., Willson, M., 1995. Class I aldolases: substrate specificity, mechanism, inhibitors and structural aspects. Progress in Biophysics and Molecular Biology 63, 301–340.

Gόmez-Arreaza, A., Acosta, H., Quiñones, W., Concepción, J.L, Michels, P.A., Avilán, L., 2014. Extracellular functions of glycolytic enzymes of parasites: Unpredicted use of ancient proteins. Molecular and Biochemical Parasitology 193, 75–81.

Guillou, F., Roger, E., Moné, Y., Rognon, A., Grunau, C., Théron, A., Mitta, G., Coustau, C., Gourbal, B.E., 2007. Excretory–secretory proteome of larval *Schistosoma mansoni* and *Echinostoma caproni*, two parasites of *Biomphalaria glabrata*. Molecular and Biochemical Parasitology 155, 45–56.

Hu, Q., Xie, H., Zhu, S., Liao, D., Zhan, T., Liu, D., 2015. Cloning, expression and partial characterization of FBPA from *Schistosoma japonicum*, a molecule on that the fluke may develop nutrition competition and immune evasion from human. Parasitology Research 114, 3459–3468.

Inoue, T., Yatsuki, H., Kusakabe, T., Joh, K., Takasaki, Y., Nikoh, N., Miyata, T., Hori, K., 1997. *Caenorhabditis elegans* has two isozymic forms, CE-1 and CE-2, of fructose-1,6-bisphosphate aldolase which are encoded by different genes. Archives of Biochemistry and Biophysics 339, 226–234.

Kochman, M., Kwiatkowska, D., 1972. Purification and properties of fructose diphosphate aldolase from *Ascaris suum* muscle. Archives of Biochemistry and Biophysics 152, 856–868.

Kovaleva, E.S., Masler, E.P., Subbotin, S.A., Chitwood, D.J., 2005. Molecular characterization of aldolase from *Heterodera glycines* and *Globodera rostochiensis*. Journal of Nematology 37, 292–296.

Laskowski, R.A., Rullmann, J.A.C., MacArthur, M.W., Kaptein, R., Thornton, J.M., 1996. AQUA and PROCHECK-NMR: programs for checking the quality of protein structures solved by NMR, Journal of Biomolecular NMR 8, 477–486. Mabiala-Bassiloua, C.G., Arthus-Cartier, G., Hannaert, V., Thérisod, H., Sygusch, J., Thérisod, M. 2011. Mannitol bis-phosphate based inhibitors of fructose 1,6-bisphosphate aldolases. ACS medicinal chemistry letters 2, 804–808.

Lebherz, H.G., Rutter, W.J., 1969. Distribution of fructose diphosphate aldolase variants in biological systems. Biochemistry 8, 109–121.

Li, S., Bian, M., Wang, X., Chen, X., Xie, Z., Sun, H., Jia, F., Liang, P., Zhou, C., He, L., Mao, Q., Huang, B., Liang, C., Wu, Z., Li, X., Xu, J., Huang, Y., Yu, X., 2014. Molecular and biochemical characterizations of three fructose-1,6-bisphosphate aldolases from *Clonorchis sinensis*. Molecular and Biochemical Parasitology 194, 36–43.

Lorenzatto, K.R., Monteiro, K.M., Paredes, R., Paludo, G.P., da Fonsêca, M.M., Galanti, N., Zaha, A., Ferreira, H.B., 2012. Fructose-bisphosphate aldolase and enolase from *Echinococcus granulosus*: genes, expression patterns and protein interactions of two potential moonlighting proteins. Gene 506, 76–84.

Marques, H.H., Zouain, C.S., Torres, C.B.B., Oliveira, J.S., Alves, J.B., Goes, A.M., 2008. Protective effect and granuloma down-modulation promoted by RP44 antigen a fructose 1,6-bisphosphate aldolase of Schistosoma mansoni. Immunobiology 213, 437–446.

Marsh, K., Lebherz, H.G., 1992. Fructose-bisphosphate aldolases; an evolutionary history. Trends in Biochemical Sciences 17, 110–118.

McCarthy, J.S., Wieseman, M., Tropea, J., Kaslow, D., Abraham, D., Lustigman, S., Tuan, R., Guderian, R.H., Nutman, T.B., 2002. *Onchocerca volvulus*: Glycolytic enzyme fructose-1,6-bisphosphate aldolase as a target for a protective immune response in humans. Infection and Immunology 20, 851–858.

Morassutti, A.L., Levert, K., Pinto, P.M., da Silva, A.J., Wilkins, P., Graeff-Teixeira, C., 2012. Characterization of *Angiostrongylus cantonensis* excretory–secretory proteins as potential diagnostic targets. Experimental Parasitology 130, 26–31.

Peng, S.-Y., Tsaihong, J.C., Fan, P.-C., Lee, K.M., 2009. Diagnosis of schistosomiasis using recombinant fructose-1,6-bisphosphate aldolase from a Formosan strain of *Schistosoma japonicum*. Journal of Helminthology 83, 211–218.

Peng, S.Y., Lee, K.M., Tsaihong, J.C., Cheng, P.C., Fan, P.C., 2008. Evaluation of recombinant fructose-1,6-bisphosphate aldolase ELISA test for the diagnosis of *Schistosoma japonicum* in water buffaloes. Research in Veterinary Science 85, 527–533.

Prompipak, J., Senawong, T., Jokchaiyaphum, K., Siriwes, K., Nuchadomrong, S., Laha, T., Sripa, B., Senawong, G., 2017. Characterization and localization of *Opisthorchis viverrini* fructose-1,6-bisphosphate aldolase. Parasitology International 66, 413–418.

Ramajo-Hernández, A., Sánchez, R.P., Martín, V.R., Oleaga, A., 2007. *Schistosoma bovis*: plasminogen binding in adults and the identification of plasminogen-binding proteins from the worm tegument. Experimental Parasitology 115, 83–91.

Reznick, A.Z., Gershon, D., 1977. Purification of fructose-1,6-diphosphate aldolase from the free-living nematode *Turbatrix aceti*. Comparison of properties with those of other class I aldolases. International Journal of Biochemistry 8, 53–59.

Reznick, A.Z., Gershon, D., 1977. Age related alterations in purified fructose-1.6-sdiphosphate aldolase from the nematode *Turbatrix aceti*. Mechanisms of Aging and Development 6, 53–59.

Rhodes, M.B., 1972. *Haemonchus contortus*: Enzymes. II. Fructose diphosphate aldolase. Experimental Parasitology 31, 332–340.

Roslan, H.A., Hossain, M.A., Gerunsin, J., 2017. Molecular and 3D-structural characterization of fructose-1,6-bisphosphate aldolase derived from *Metroxlon sagu*. Brazilian Archives of Biology and Technology 60, 1–21.

Sánchez, L.B., Horner, D.S., Moore, D.V., Henze, K., Embley, T., Müller, M., 2002. Fructose-1,6-bisphosphate aldolases in amitochondriate protists constitute a single protein subfamily with eubacterial relationships. Gene 295, 51–59.

Sassi, A.J., Geary, J.F., Leroux, L.P., Moorhead, A.R., Satti, M., Mackenzie, C.D., Geary, T.G., 2014. Identification of *Dirofilaria immitis* proteins recognized by antibodies from infected dogs. Journal for Parasitology 100, 364–367.

Schrodinger, L.L.C., 2010. The PyMOL molecular graphics system. Version 1, 0.

Sun, Z., Shen, B., Wu, H., Zhou, X., Wang, Q., Xiao, J., Zhang, Y., 2015. The secreted fructose-1,6-bisphosphate aldolase as a broad-spectrum vaccine candidate against pathogenic bacteria in aquaculture. Fish and Shellfish Immunology 46, 638–647.

Tittmann, K., 2014. Sweet siblings with different faces: The mechanisms of FBP and F6P aldolase, transaldolase, transketolase and phosphoketolase revisited in light of recent structural data. Bioorganic Chemistry 57, 263–280.

Umair, S., Ria, C., Knight, J.S., Simpson, H.V., 2013. Sarcosine metabolism in *Haemonchus contortus* and *Teladorsagia circumcincta*. Experimental Parasitology 134, 1–6.

Wiederstein, M., Sippl, M.J., 2007. ProSA-web: interactive web service for the recognition of errors in three-dimensional structures of proteins, Nucleic Acids Research 35, W407–W410.

Yang, J., Yan, R., Roy, A., Xu, D., Poisson, J., Zhang, Y., 2015. The I-TASSER Suite: Protein structure and function prediction. Nature Methods 12, 7–8.

Yan, R., Xu, L., Wang, J., Xiaokai, S., Xiangrui, L., 2013. Cloning and characterization of aldolase from parasitic nematode *Haemonchus contortus*. Journal of Animal and Veterinary Advances 12, 478–486.

Zhang, Y., Skolnick, J. 2004. Scoring function for automated assessment of protein structure template quality, Proteins 57, 702–710.

Zeelon, P., Gershon, H., Gershon, D., 1973. Inactive enzyme molecules in aging organisms. Nematode fructose 1, 6-diphosphate aldolase. Biochemistry 12, 1743–1750.

